# C1q limits cystoid edema by maintaining basal beta-catenin-dependent signaling and blood-retina barrier function

**DOI:** 10.1101/2025.07.22.666172

**Authors:** Lingling Zhang, Jacklyn Levey, Md. Abedin, Ha-Neul Jo, Emmanuel Odame, Kaia Douglas, Elise Thoreen, Scott W. McPherson, Heidi Roehrich, Somasekar Seshagiri, Stephane Angers, Zhe Chen, Harald J. Junge

## Abstract

Macular edema (ME) causes significant vision impairment and occurs in several prevalent retinal diseases including diabetic retinopathy (DR), choroidal neovascularization (CNV), retinal vein occlusion, and uveitis. Retinal edema typically results from dysfunction of the blood-retina barrier (BRB), which is associated with increased retinal expression of complement components. It is unclear whether the classical complement pathway has detrimental or protective roles in the context of BRB dysfunction. Here, we characterize *Tspan12* KO^DBM^ (<u>D</u>isrupted <u>B</u>arrier <u>M</u>aintenance) mice, a new mouse model of cystoid edema based on genetically and pharmacologically manipulating beta-catenin-dependent norrin/frizzled4 (FZD4) signaling. We assess BRB function, cystoid edema, ERG, and microglia activation outcomes in an aging study with WT, *C1qa* KO, *Tspan12* KO^DBM^, and *Tspan12* KO^DBM^*; C1qa* KO compound mutant mice. Phenotypic analyses and cell-based experiments indicate that C1QA contributes to maintaining basal beta-catenin-dependent signaling and that the absence of C1QA exacerbates BRB dysfunction, cystoid edema, and neuroinflammation in *Tspan12* KO^DBM^*; C1qa* compound mutant mice. Activation of beta-catenin-dependent signaling by a FZD4/LRP5 agonist antibody modality achieves complete resolution of cystoid edema. This study shows that reducing or enhancing norrin/frizzled4 signaling can increase or decrease cystoid edema, respectively, underscoring its potential as a therapeutic target in ME. Furthermore, this study provides novel insights into the contribution of C1QA to BRB maintenance.

## Introduction

Macular edema (ME) arises when the rate of fluid entry into the retina significantly exceeds the rate of fluid export, resulting in a pathological accumulation of fluid. The macula is prone to the formation of edema, likely due to its anatomic specialization. Edema may appear as diffuse fluid accumulation associated with increased macula thickness, fluid accumulation in the subretinal space, or fluid in intraretinal cyst-like spaces (cystoid edema, CE). ME poses a significant risk to acute central vision and may lead to irreversible neural damage. This condition often develops as a complication of prevalent eye diseases, including diabetic retinopathy, age-related macular degeneration, retinal vein occlusion, and uveitis (1).

A common cause of ME is dysfunction of the inner BRB, where vascular endothelial cells (ECs) play a pivotal role as a critical cellular component. The BRB plays a key role in maintaining the retinal microenvironment by regulating transport at the interface of the circulatory system and the neural retina. ECs of an intact BRB are characterized by efficient tight junctions, low rates of transcytosis, close interactions with glia cells, a high degree of pericyte coverage, and the expression of transporters for nutrients, hormones, and metabolic waste (2). With an intact BRB, oncotic pressure (driving water flow into blood vessels towards the high concentration of plasma proteins) and hydrostatic pressure (driving water flow out of blood vessels) provide a degree of balance so that fluid extravasation does not overwhelm retinal export mechanisms into the vitreous or across the retinal pigment epithelium. With BRB breakdown and the resulting protein extravasation, oncotic pressure is reduced and fluid accumulation in the retina increases (3). In addition, protein extravasation (e.g., fibrinogen) causes inflammation (4).

The major pathway that induces and maintains BRB function is the norrin/frizzled4 (FZD4) pathway. The secreted protein norrin (gene symbol NDP, for Norrie Disease Pseudoglioma) is released from Müller glia and horizontal cells and activates beta-catenin-dependent signaling in ECs by binding to a receptor complex containing the membrane proteins FZD4, low-density lipoprotein receptor-related 5 (LRP5), and tetraspanin 12 (TSPAN12). This signaling pathway plays a crucial role in enabling retinal angiogenesis, BRB induction, and BRB maintenance (5–7). The loss of norrin signaling in mice can cause CE, even in the absence of a macula (8), however, norrin gene disrupted mice are characterized by compounding pathologies including vascular malformations and hypoxia (9), complicating the analysis of pathological roles of BRB dysfunction in CE. TSPAN12 is a co-receptor for norrin (10, 11) and is required for norrin/frizzled4 signaling (12). *Tspan12* endothelial cell-specific knockout (ECKO) mice have been used to separate angiogenesis and BRB defects by inducing tamoxifen-induced recombination after angiogenesis is complete. While this model circumvents the occurrence of vascular malformations and hypoxia, CE in *Tspan12* ECKO mice is variable (7). Improved mouse models that display extensive CE despite the lack of a macula are needed. This is important for better understanding of CE and to test therapeutic approaches. For example, the activation of the norrin/frizzled4 pathway by agonist antibodies that bind FZD4 and LRP5 or FZD4 and LRP6 (13–15) alleviate BRB defects and suppress neovascularization in oxygen-induced retinopathy models (16–18). FZD4/LRP5 and FZD4/LRP6 agonists emerge as a novel class of agonists with potential uses in ME, but efficacy in preclinical mouse models of retinal edema remains to be demonstrated.

We previously reported that complement components are elevated in mice with impaired BRB maintenance (7). This finding sparked questions about the role of complement in the context of BRB dysfunction. The classical complement pathway is part of the innate immune system and is implicated in retinal disease (19). The C1 complex starts the classical complement cascade by activation of C1r and C1s serine proteases. C1q (which is composed of 6 heterotrimers, each containing one C1QA, C1QB and C1QC polypeptide) serves as essential scaffold in the C1 complex (20).

Upon encountering antigen-antibody complexes or other activators, e.g., phosphatidyl serine on apoptotic cells (21) or pentraxins (22), the C1 complex initiates a cascade that can lead to cell destruction via the membrane attack complex, or cell opsonization and phagocytosis (23). C1q binds to multiple types of complement receptors to promote phagocytosis (24).

The classical complement system is important for maintaining tissue homeostasis, e.g., in synapse elimination (25) and wound healing (26). Pathophysiological actions include roles in neurodegenerative disease (27, 28), glaucoma (29), and in autoimmune diseases (30). Among the latter, C1q deficiency is a major risk factor for systemic lupus erythematosus (SLE) (31), as insufficient clearance of apoptotic cells and nuclear antigens promotes autoantibody generation in SLE (32). Immunoglobulin extravasation in the retina due to BRB dysfunction could further promote autoimmunity in C1q deficiency. Whether *C1qa* KO mice develop retinal manifestations of SLE (including hemorrhages, and cotton-wool spots) when C1q deficiency is compounded with strong BRB defects and immunoglobulin extravasation, is not known.

The classical complement pathway is implicated in retinal diseases characterized by BRB dysfunction (19). Furthermore, increased retinal complement component expression was observed in mice with BRB dysfunction (7). This raises the question if BRB dysfunction could drive retinal disease progression via the classical complement pathway. Abundant extravasated immunoglobulin could form antigen-antibody complexes on the cell surface of retinal cells that activate the classical complement pathway, which may enhance phagocytosis and cause retinal damage. However, protective roles for C1q are also plausible. C1-dependent cleavage of LRP6 and activation of beta-catenin-dependent signaling was reported in skeletal muscle regeneration and arterial remodeling (33, 34). Therefore, C1q could be required for BRB maintenance by promoting basal beta-catenin-dependent signaling in ECs. Together, whether the classical complement pathway in the context of BRB dysfunction has detrimental or protective roles, remains poorly understood.

Here, we establish a new mouse model of BRB maintenance defects and CE based on genetically and pharmacologically manipulating norrin/frizzled4 signaling at specific stages of retinal vascular development. We use this model to test the role of the classical complement system in the context of BRB breakdown. Interestingly, we find that the loss of C1QA exacerbates BRB dysfunction and associated pathologies.

Results from cell-based studies indicate a role of C1q in maintaining basal levels of beta-catenin dependent signaling in ECs, revealing a novel role of C1q in BRB maintenance. Furthermore, our analysis shows that increasing or decreasing norrin/frizzled4 signaling modulates CE. We find that FZD4/LRP5 agonists alleviate BRB dysfunction and completely resolve CE, highlighting that norrin/frizzled4 signaling is a highly suitable target for pharmacological intervention in ME.

## Results

### *Tspan12* KO^DBM^ mice display extensive cystoid edema

*Tspan12* ECKO mice develop and maintain a normal deep vascular plexus if tamoxifen-dependent recombination is induced after the phase of retinal developmental angiogenesis is over. While the three-layered retinal vasculature is maintained, *Tspan12* ECKO mice exhibit BRB defects and moderate CE due to impaired norrin/frizzled4 signaling (7). We found that this model was useful to correlate defined leakage areas directly with sites of CE formation using fluorescein angiography (FA) guided optical coherence tomography (OCT) (Fig. 1A). However, Cre-mediated recombination in *Tspan12* ECKO mice is not complete (18), resulting in only mild or moderate CE that is not fully penetrant (Fig. 1A and D). To generate a model with increased and more penetrant CE, we sought to use a model with complete *Tspan12* gene inactivation while bypassing vascular malformations and hypoxia, which are characteristic phenotypes of *Tspan12* KO mice. We used a strategy of pharmacologically and genetically manipulating norrin/frizzled4 signaling at specific developmental stages to achieve that. Angiogenesis and BRB phenotypes of *Tspan12* KO mice were rescued by administration of F4L5.13, a FZD4/LRP5 agonist antibody and norrin mimetic, every three days from P6 to P28, as described previously (18). The administration of F4L5.13 during postnatal development supports virtually normal retinal angiogenesis and formation of the BRB in *Tspan12* KO mice (18). However, after cessation of treatment, norrin/frizzled4 signaling is no longer activated and therefore the maintenance of the BRB is disrupted (schematic in Fig. 1B). Hereafter, we refer to this new model as *Tspan12* KO^DBM^, for <u>D</u>isrupted <u>B</u>RB <u>M</u>aintenance. 6 months after cessation of treatment, *Tspan12* KO^DBM^ mice displayed severe BRB dysfunction and widespread CE (Fig. 1C). We assigned each retina a cystoid edema score ranging from 0 to 5 using a grading scale described in Supplemental Fig. S1. This analysis revealed that the vast majority of *Tspan12* KO^DBM^ mice displayed CE, whereas CE in *Tspan12* ECKO was sporadic and milder (Fig. 1D). CE in both models was predominantly detected in the inner nuclear layer, correlating with the high density of leaky intraretinal capillaries that flank the inner nuclear layer on both sides. Together, by genetic and pharmacological manipulation of norrin/frizzled4 signaling, we created a mouse model with extensive CE.

**Fig. 1.**
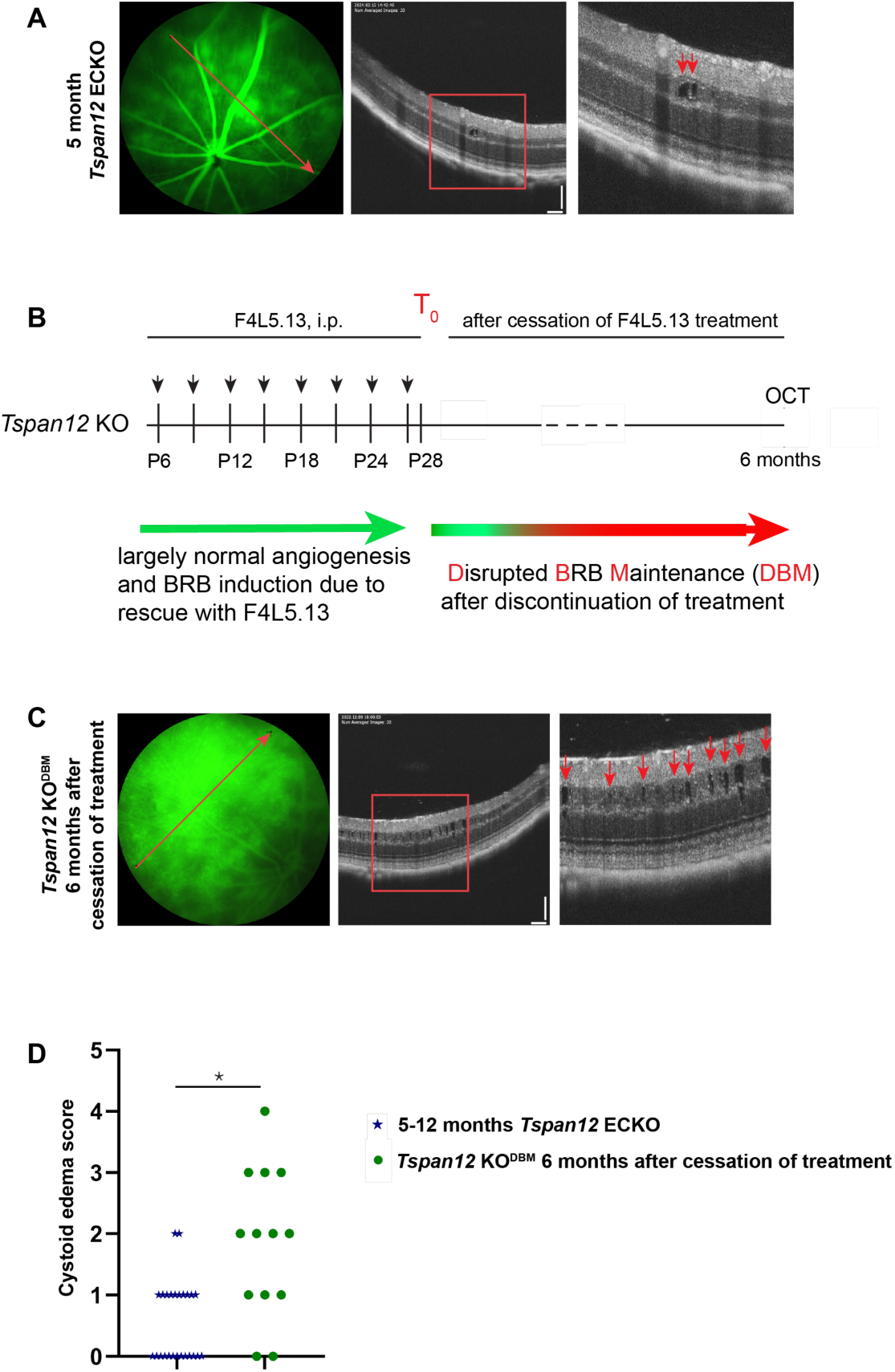
Cystoid edema is more severe in *Tspan12* KO^DBM^ mice compared to *Tspan12* ECKO mice. (A) Representative fluorescein angiography fundus images and OCT scan images show *Tspan12* ECKO mice with spotty retinal vascular leakage and moderate CE lesions. The red lines show the OCT line scan relative to the FA image. Boxed areas are shown enlarged in the panels on the right. Red arrows point to CE lesions. Scale bars: 100 µm. (B) Schematic representation of F4L5.13 administration to *Tspan12* KO mice until P28 (T_0_) and subsequent cessation of treatment to generate *Tspan12* KO^DBM^ (“<u>D</u>isrupted <u>B</u>RB <u>M</u>aintenance”) mice. (C) Representative fluorescein angiography fundus images and OCT scan images show *Tspan12* KO^DBM^ mice with extensive CE. Scale bars: 100 µm. (D) Cystoid edema scores in *Tspan12* ECKO mice, and *Tspan12* KO^DBM^ mice, n=13 *Tspan12*^DBM^ and n=21 *Tspan12* ECKO retinas, a Mann-Whitney non-parametric test was used to test for differences of data on an ordinal scale.

### *Tspan12* KO^DBM^; *C1qa* KO compound mutant mice display increased retinal vascular leakage and cystoid edema compared to *Tspan12* KO^DBM^ mice

Next, we use this model of CE to better define the role of the classical complement system in the context of BRB dysfunction. To inactivate the classical complement pathway, we used a *C1qa* null allele (35), as C1QA is a structural component of the C1 complex necessary for the activation of the classical complement cascade (20). Four groups of mice were followed in a year-long longitudinal FA study to assess BRB function: WT, *C1qa* KO, *Tspan12* KO^DBM^, and *Tspan12* KO^DBM^; *C1qa* KO compound mutant mice. *Tspan12* KO^DBM^ were generated by repeated administration of F4L5.13 until P28, which was defined as timepoint T_0_ (Fig. 2A). FA at P28 confirmed our previous report that F4L5.13 rescues the BRB phenotypes of *Tspan12* KO mice (18). At T_0_ + 2 weeks, the BRB was already substantially compromised due to cessation of treatment. There was no notable leakage in the retinas of *C1qa* single KO mice, which appeared comparable to WT control. *Tspan12* KO^DBM^; *C1qa* KO compound mutant mice exhibited increased leakage compared to *Tspan12* KO^DBM^ mice, however, non-quantitative FA was not sufficient to ascertain this difference (Fig. 2B). Therefore, we quantified sulfo-NHS-LC-biotin leakage (a terminal procedure) in all 4 groups at the T_0_ + 12 months end point of the study.

**Fig. 2.**
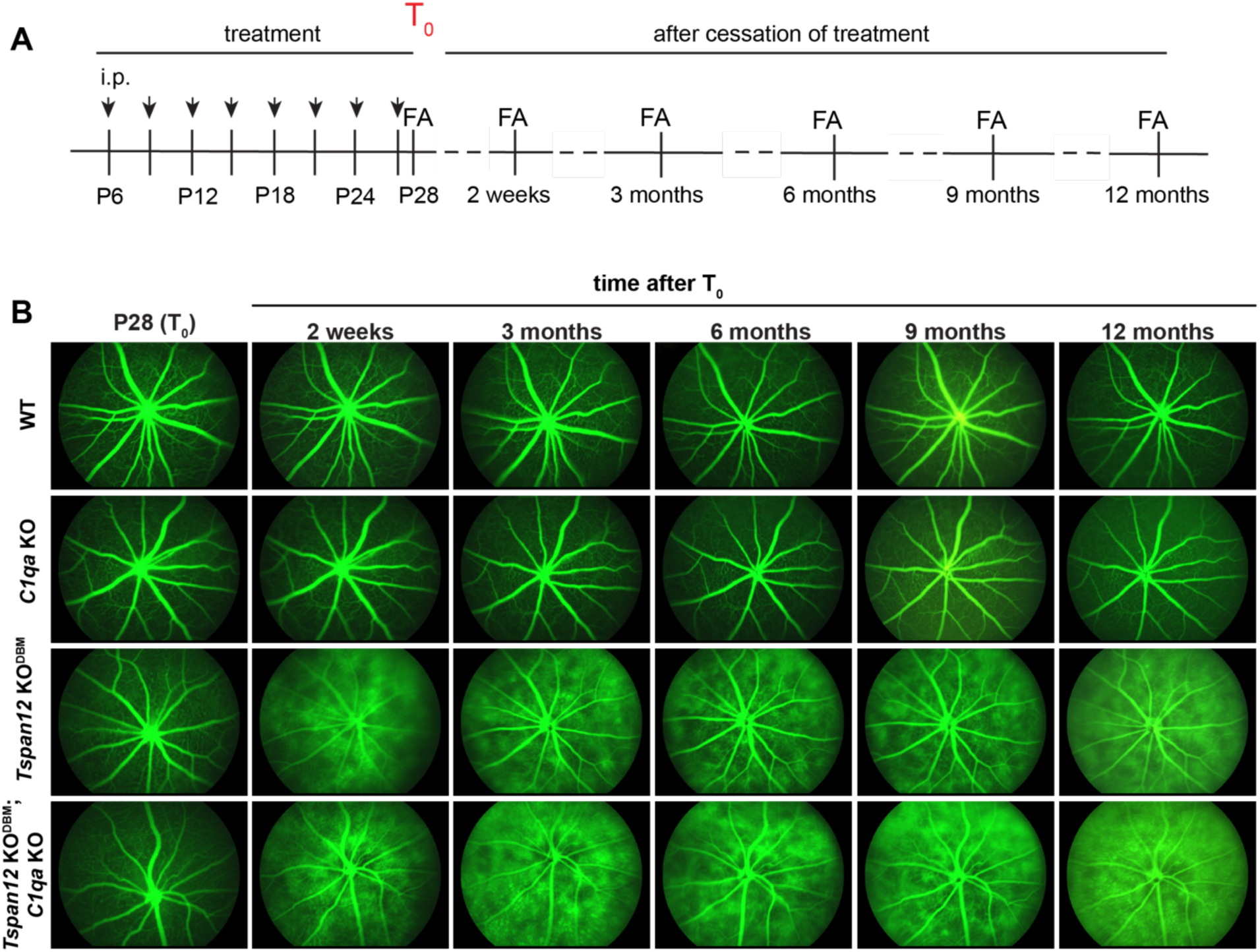
Longitudinal FA imaging in four groups of mice. (A) Schematic overview of the longitudinal study design. (B) The images are representative of 5-8 mice per group with similar results.

Sulfo-NHS-LC-biotin is a reactive tracer that biotinylates plasma proteins and luminal retinal EC proteins while it circulates through the vasculature. When the BRB is impaired, the tracer leaks out of retinal blood vessels and biotinylates proteins in the vascular basement membrane, perivascular space, and retinal parenchyma. Covalently attached biotin is subsequently detected using fluorescent streptavidin probes. Unlike diffusible FITC, biotin signal is not attenuated during wash steps and is therefore less variable. We used this approach to quantify BRB leakage in 12-month-old mice from retinal whole mounts, which were imaged using parameters optimized for the strong signal in *Tspan12* KO^DBM^ mice and *Tspan12* KO^DBM^; *C1qa* KO compound mutant mice. This experiment revealed a moderate but statistically significant increase of tracer extravasation in *Tspan12* KO^DBM^; *C1qa* KO compound mutant mice compared to *Tspan12* KO^DBM^ mice (Figure 3A and B).

**Fig. 3.**
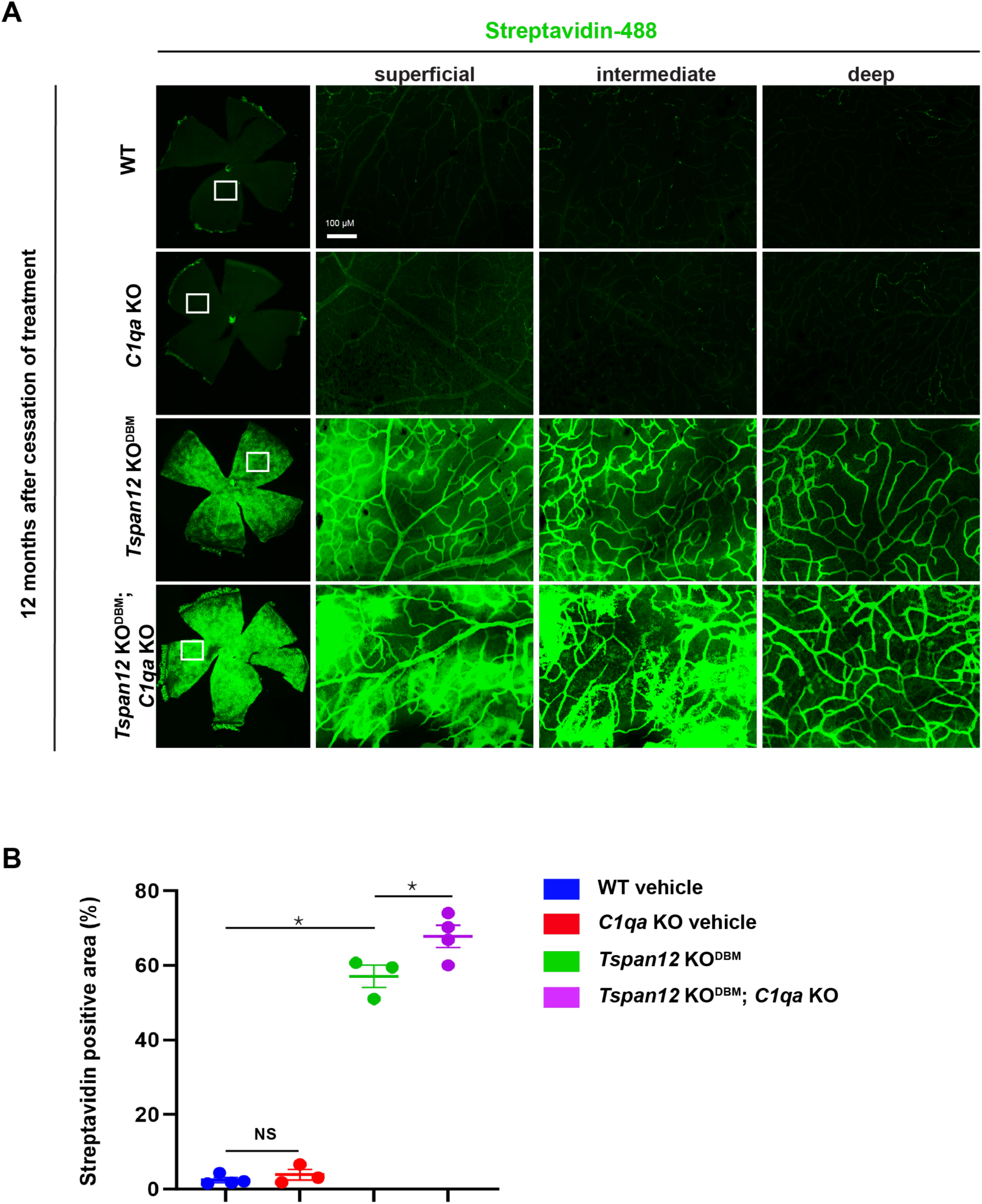
Increased sulfo-NHS-LC-biotin tracer leakage in *Tspan12* KO^DBM^; *C1qa* KO compound mutant retinas compared to *Tspan12* KO^DBM^ retinas. (A) Left panels: Stitched 4x images of flat mount retinas. Right panels: 20x projections of the areas demarcated by white boxes. 3D image stacks were acquired and projections for each of three vascular layers were generated. Image acquisition settings were optimized for the strong signal in *Tspan12* KO^DBM^ mice. Scale bar: 100 µm. (B) Streptavidin positive area above threshold was quantified. 4 areas per retina were imaged and the data from each retina was averaged. N = 3-4 retinas per group. Average +/-SEM is shown. Asterisks indicate significance using a one-way ANOVA with Tukey post hoc.

6-9 months after cessation of treatment with F4L5.13, significant BRB leakage and CE were observed in the retinas of *Tspan12* KO^DBM^ mice *and Tspan12* KO^DBM^; *C1qa* KO compound mutant mice. In most compound mutant mice CE appeared more severe than in single mutant *Tspan12* KO^DBM^ mice (Fig. 4A). Grading the CE severity revealed that at T_0_ + 6 months, CE scores in compound mutant mice trended to be higher than in single mutant mice but did not quite reach significance (P=0.0507, Fig. 4B). At T_0_ + 9 months, CE scores of compound mutant mice were significantly higher than those of single mutant mice (Fig. 4C). Thus, the severity of CE scores correlated with the increased severity of sulfo-NHS-LC-biotin leakage in *Tspan12* KO^DBM^; *C1qa* KO compound mutant mice compared to *Tspan12* KO^DBM^ single mutant mice.

**Figure 4.**
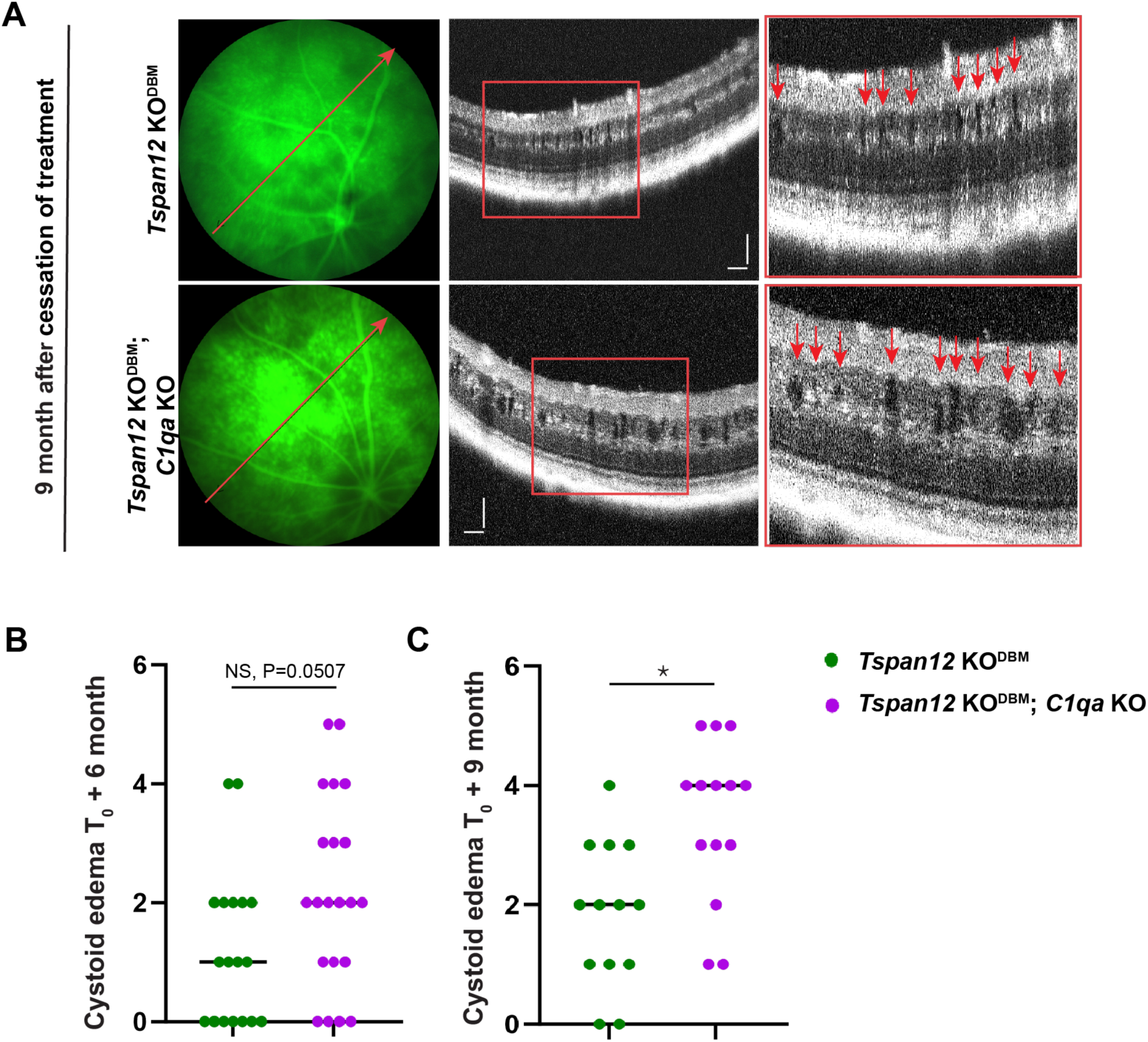
Increased cystoid edema in *Tspan12 KO^DBM^*; *C1qa* KO compound mutant retinas compared to *Tspan12 KO^DBM^*retinas. (A) Representative fluorescein angiography fundus images and OCT scan images. The red lines show the OCT line scan relative to the FA image. Boxed areas are shown enlarged in the panels on the right. Red arrows point to CE lesions. Scale bars: 100 µm. (B) At T_0_ + 6 months, cystoid edema scores in the *Tspan12* KO; *C1qa* KO mice trended to be higher than in the *Tspan12* KO mice but did not reach significance. N=18-21 retinas, a Mann-Whitney non-parametric test was used as the CE scores are on an ordinal scale. (C) At T_0_ + 9 months, a significant difference in CE scores was observed. N=13-14 retinas, Mann-Whitney non-parametric test.

### ERG b-wave defects in *Tspan12* KO^DBM^ and *Tspan12* KO^DBM^; *C1qa* KO mice

We used dark-adapted electroretinography to examine retinal functional differences at three time points: T_0_, T_0_ + 3 month and T_0_ + 10 month. As expected, ERG a-wave and b-wave amplitudes in all groups declined with age (36). The b-wave reflects the net effect of ion currents across the membranes of inner retinal cell populations, predominantly bipolar cells, in response to rod activity. The a-wave reflects the net effect of ion currents across the membranes of the rod cell population. At T_0_, the b-wave amplitude of *Tspan12* KO^DBM^ mice compared to WT mice was not significantly changed, because repeated administration of F4L5.13 rescues postnatal retinal angiogenesis and restores the scotopic ERG b-wave in *Tspan12* KO mice (18). However, a striking decrease of the b-wave amplitude of *Tspan12* KO^DBM^ and *Tspan12 KO^DBM^*; *C1qa* KO compound mutant mice was observed at the T_0_ + 3 months and T_0_ + 10 months time points compared to both control genotypes (Fig. 5A). This strong reduction of the ERG b-wave correlated with the strong BRB defects and CE in *Tspan12* single and compound mutant mice. In contrast, the dark-adapted a-wave was not significantly changed (Fig. 5B). Accordingly, the b/a amplitude ratio (expressed as absolute value) was significantly lower in *Tspan12* KO^DBM^ and *Tspan12* KO^DBM^; *C1qa* KO compound mutant mice at T_0_ + 3 months and T_0_ + 10 months compared to both control genotypes (Fig. 5C). While the b/a ratio trended to be smaller in *Tspan12* KO^DBM^; *C1qa* KO compound mutant mice compared to *Tspan12* KO^DBM^ single mutant mice at early time points, this difference became significant at the T_0_ + 10 months time point, consistent with the increased sulfo-NHS-LC-biotin leakage and CE in the compound mutant group. In addition, we observed a significant increase of the b/a ratio in *C1qa* KO mice compared to WT mice at T_0_ and a non-significant trend towards higher b/a ratios at the later time points, which may reflect developmental differences at the photoreceptor triad synapse in *C1qa* KO mice (37). We note that increased b/a ratios due to loss of C1QA may lead to an underestimation of the reduction of b/a ratios in *Tspan12 KO^DBM^*; *C1qa* KO compound mutant mice compared to *Tspan12* KO^DBM^ single KO mice.

**Fig. 5.**
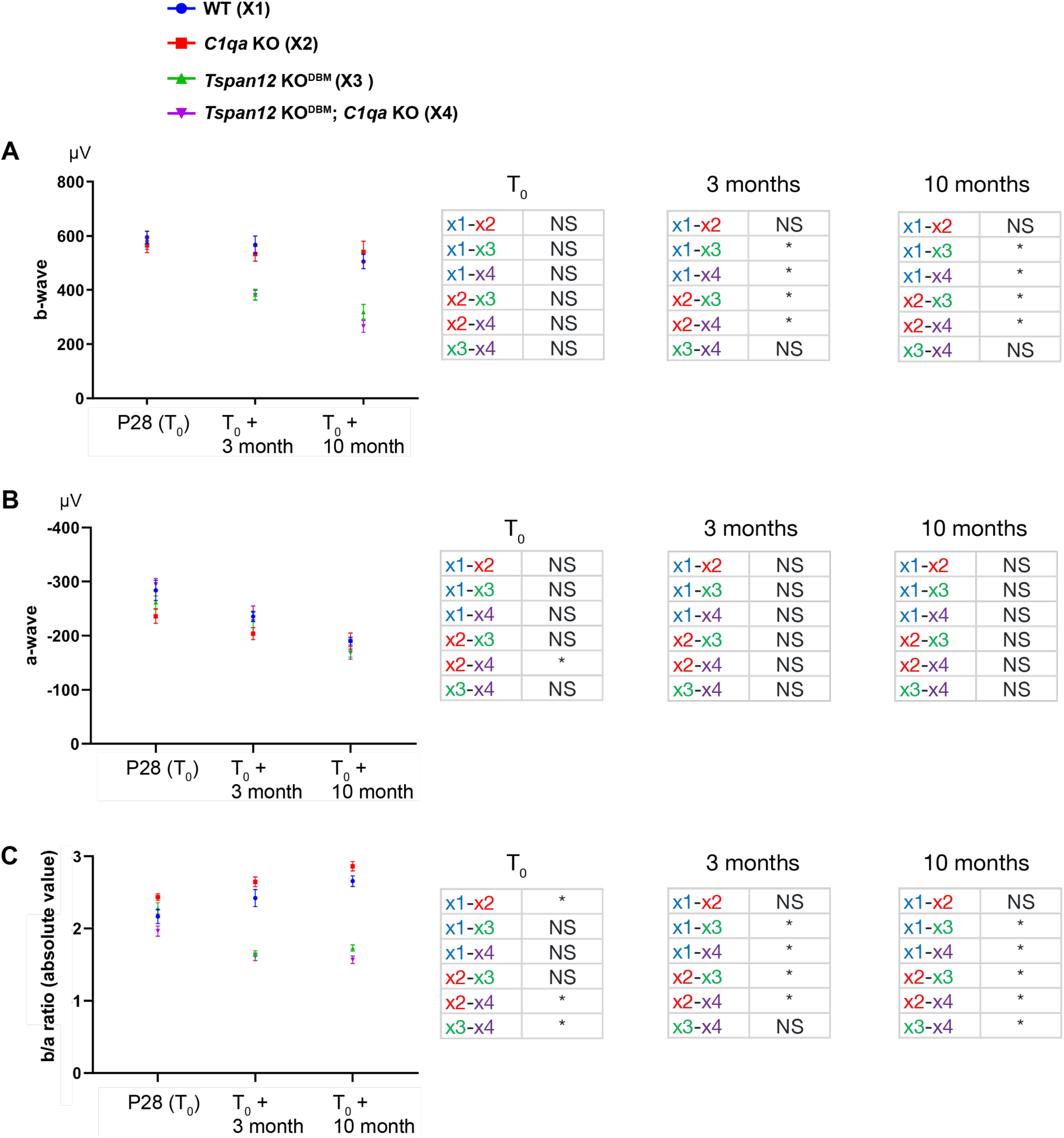
ERG b-wave reduction in mice with BRB maintenance defects. (A) P28: n=16-24 retinas. T_0_ +3 month: n=16-24 retinas. T_0_ +10 month: n=14-16 retinas. One-way ANOVA with Tukey post hoc, error bars represent SEM. At the T_0_ +3 months time point, a non-parametric Kruskal-Wallis test was used due to non-equal variances. (B) ERG a-wave of the same mice as detailed above. One-way ANOVA with Tukey post hoc, average +/- SEM shown. (C) b/a ratio of the same mice as detailed above. One-way ANOVA with Tukey post hoc, average +/- SEM shown. At the T_0_ +3 months time point, a non-parametric Kruskal-Wallis test was used due to non-equal variances.

### Microglia and macroglia phenotypes in *Tspan12* KO^DBM^ and *Tspan12* KO^DBM^; *C1qa* KO mice

Endothelial blood-CNS barrier breakdown triggers neuroinflammation, for example due to the extravasation of fibrinogen (4). Conversely, neuroinflammation is a cause for barrier dysfunction. This reciprocal relationship can lead to amplification of both barrier dysfunction and neuroinflammation (38). Expansion of microglia or infiltration of monocyte-derived macrophages can be a feature of neuroinflammation. We stained for IBA-1, a marker of resident retinal microglia and monocyte-derived macrophages, after tissue was harvested at the T_0_ + 12-month endpoint of the study. We found a strong IBA1^+^ microglia/macrophage cell expansion in *Tspan12* KO^DBM^ and *Tspan12* KO^DBM^; *C1qa* KO compound mutant mice (Fig. 6A). Expansion of IBA-1^+^ cells was more severe in the compound mutant mice (Fig. 6B). Consistent with this quantification, we found a significant increase of the mRNA *Trem2*, a marker for microglia/monocyte-derived macrophages. *Trem2* was increased in *Tspan12* KO^DBM^ and *Tspan12* KO^DBM^; *C1qa* KO compound mutant mice, this increase trended to be more pronounced in *Tspan12* KO^DBM^; *C1qa* KO compound mutant mice compared to *Tspan12* KO^DBM^ single mutant mice, although the difference did not reach significance (P = 0.11) (Fig. 6C).

**Fig. 6.**
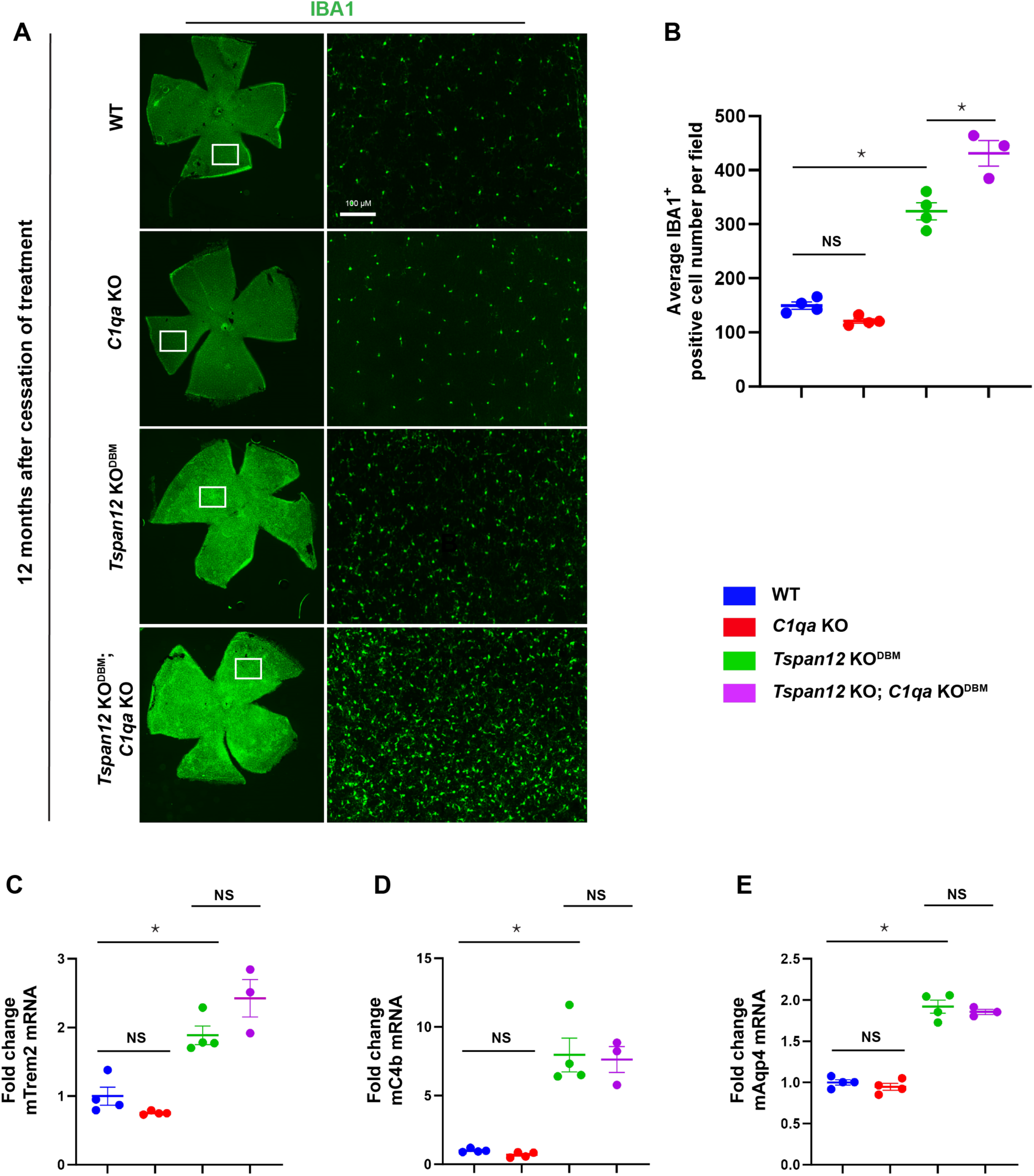
Microglia and macroglia phenotypes in mice with BRB maintenance defects. (A) 4x stitched images of whole-mount retinas stained for the microglia/monocyte-derived macrophage marker IBA1. Boxed areas are imaged at 20x and shown in the right panel. Scale bar: 100 µm. (B) Quantification of IBA1^+^ cells per 20 x field of view. N=3-4 retinas per group, one-way ANOVA with Tukey post hoc, average +/- SEM shown. (C-E) RT-qPCR for the indicated genes, N=3-4 retinas per group, ANOVA with Tukey post hoc, average +/- SEM shown.

How glia responds to endothelial blood-CNS barrier dysfunction is incompletely understood. We found that the mRNAs of two retinal glia markers, *C4b (*Fig. 6D) and *Aqp4* (aquaporin 4) (Fig. 6E) were upregulated in *Tspan12* KO^DBM^ and *Tspan12* KO^DBM^; *C1qa* KO compound mutant mice. C4b may help in the clearance of extravasated protein or cell debris via opsonization (39) and is expressed in retinal astrocytes and activated microglia (9). AQP4, which is expressed in Müller cells and astrocytes, is involved in water export from the retina (3) and may partially compensate for the increased fluid entry through a leaky BRB.

Although C1q deficiency can lead to SLE with retinal manifestations in human patients, e.g., hemorrhages, and microinfarctions in the nerve fiber layer (cotton-wool spots) (40), such phenotypes have not been reported in the retinas of *C1qa* KO mice. Because immunoglobulin extravasation in *Tspan12* KO^DBM^; *C1qa* KO compound mutant mice may promote autoimmunity, we examined the fundus of mice of all four genotypes. However, immunoglobulin extravasation and C1q deficiency was not sufficient to cause hemorrhages, or cotton-wool spots (Supplemental Fig. S2) on a C57BL/6J background (see discussion).

### Loss of C1QA dampens ligand-independent basal beta-catenin-dependent signaling

In skeletal muscle regeneration and arterial remodeling, complement C1q promotes basal beta-catenin-dependent signaling in a wnt-independent manner by binding to frizzled receptors and inducing C1s-dependent cleavage of the ectodomain of the wnt co-receptor LRP6 (33, 34). We wondered if loss of C1QA in *Tspan12* KO^DBM^; *C1qa* KO compound mutant mice reduces basal beta-catenin signaling in ECs in a context where ligand-induced norrin/frizzled4 signaling is already strongly impaired. A reduction of basal signaling could explain the exacerbated BRB leakage and increased CE in *Tspan12* KO^DBM^; *C1qa* KO compound mutant mice compared to *Tspan12* KO^DBM^ mice. To test this hypothesis, we performed TOPFlash luciferase reporter assays using 293T cells transfected with FZD4 and LRP5. Cells were cultured in medium with serum from aged WT mice, a major source of C1Q (33), or serum from age matched *C1qa* KO mice. Basal signaling activity (without norrin stimulation) was significantly decreased in cells cultured in C1QA-deficient medium compared to medium with serum from WT mice (Fig. 7A), consistent with a previous report that used WT vs. *C1qa* KO serum to stimulate basal signaling in other biological contexts (33). Basal reporter activity was also reduced in cells co-expressing the TSPAN12 co-receptor for norrin (Fig. 7B). To rule out the possibility that norrin contributed to activating the TOPFlash reporter, we compared TOPFlash activity of cells cultured in medium with WT serum vs. serum from *Ndp* (norrie disease protein) KO mice and found that reporter activity was unchanged, confirming that serum is not a source of norrin (Fig. 7C). Norrin-induced signaling in TOPFlash reporter assays is substantially higher than basal receptor signaling (10). The relatively lower activity of basal signaling compared to ligand-induced signaling likely explains why the additional increase of BRB leakage and CE formation in *Tspan12* KO^DBM^; *C1qa* KO compound mutant mice is moderate, and why *C1qa* single KO mice (which maintain strong norrin-induced frizzled4 signaling) have no obvious BRB phenotypes. Together, our data indicate that C1Q-mediated activation of beta-catenin-dependent signaling provides basal activity that is important in the context of a dysfunctional BRB. Disease contexts in which this mechanism is likely relevant include familial exudative vitreoretinopathy (FEVR), which is a retinal vascular disease caused by impaired norrin/frizzled4 signaling, for example by mutations in the human *TSPAN12* gene (41).

**Fig. 7.**
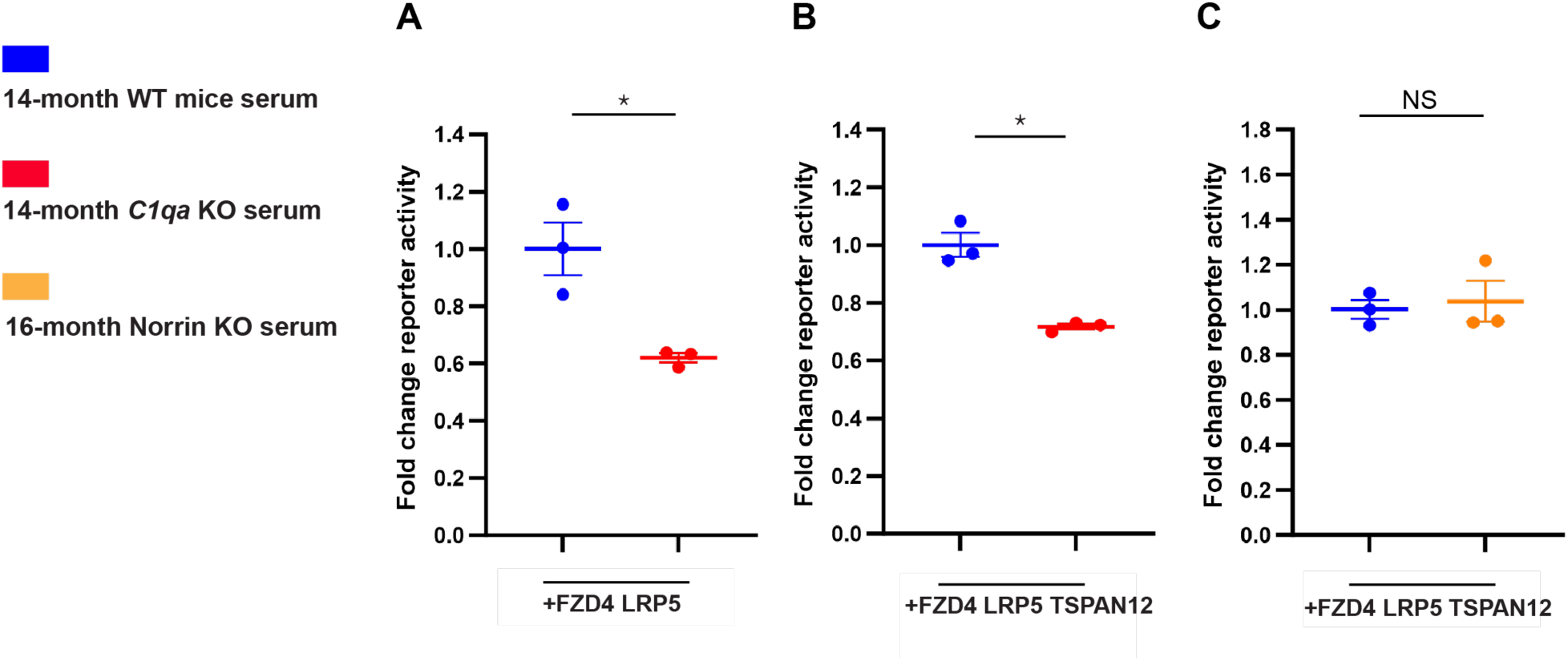
Loss of C1QA dampens ligand-independent basal beta-catenin-dependent signaling through Frizzled4. (A-C) TOPFlash Dual-Glo assay (firefly activity divided by renilla activity) in 293T cells transfected with the indicated constructs. Cells were cultured in serum from mice of different genotypes as indicated in the figure. N=3 biological replicates, average +/- SEM shown. Welch’s heteroscedastic t-test.

### F4L5.13 achieves complete resolution of cystoid edema in treatment naïve ***Tspan12* ECKO mice**

We previously reported that F4L5.13 restores BRB function in *Tspan12* ECKO mice (18), therefore, we wondered if the norrin mimetic can also alleviate CE. We used the *Tspan12* ECKO model for this experiment as these mice are treatment naïve. Three doses of F4L5.13 were administered to 5-month-old *Tspan12* ECKO mice i.p. every two days (Fig. 8A). Image-guided OCT showed moderate CE before treatment. The same mouse eye was re-imaged 11 days later. Strikingly, F4L5.13 treatment rescued BRB leakage and CE was completely resolved in all retinas (Fig. 8B and C). This finding demonstrates the efficacy of FZD4/LRP5 agonists in a mouse model of CE and reinforces the concept that the norrin/frizzled4 signaling pathway is a highly suitable target for pharmacological intervention in ME.

**Fig. 8.**
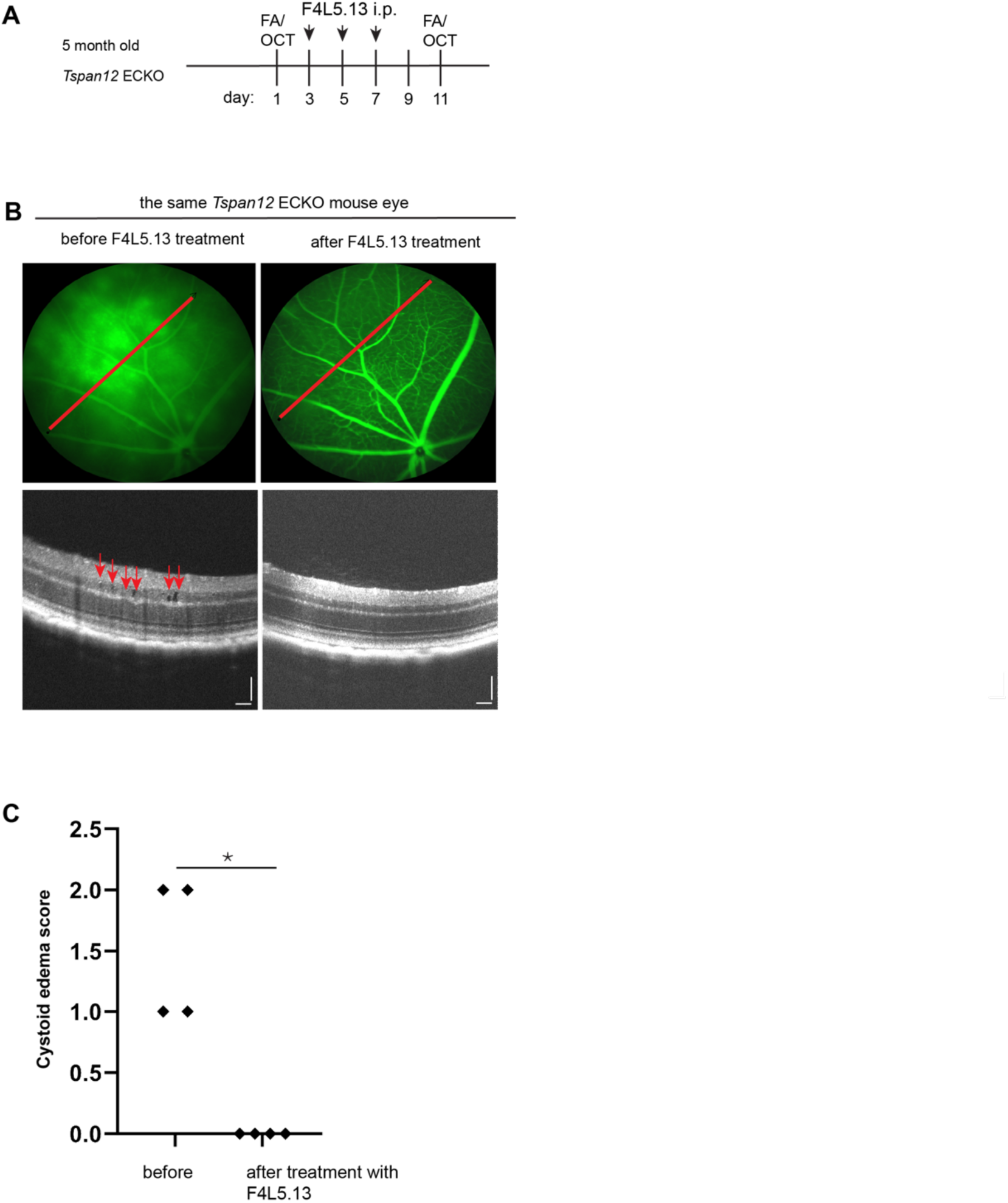
F4L5.13 achieves complete resolution of cystoid edema in treatment naïve *Tspan12* ECKO mice. (A) Schematic representation of the experimental design. Animals received three doses of F4L5.13, 10 mg/kg, i.p. (B) Representative fluorescein angiography fundus images and OCT scan images show *Tspan12* ECKO mice with retinal vascular leakage and moderate CE lesions. The red lines show the OCT line scan relative to the FA image. Right panels show the same retina after treatment. Scale bars: 100 µm. (C) A Mann-Whitney non-parametric test was used to test for differences of CE scores on an ordinal scale, n=4 retinas.

## Discussion

A major conclusion from the present study is that loss of C1QA, which is required for the initiation of the classical complement cascade, reduces basal beta-catenin-dependent signaling activity in ECs and exacerbates BRB dysfunction in a context of impaired norrin/frizzled4 signaling. This conclusion is based on phenotypic comparison of *Tspan12* KO^DBM^; *C1qa* KO compound mutant mice, cell-based assays for basal FZD4/LRP5 signaling as well as prior studies that established a relationship of C1q and basal frizzled signaling in other biological contexts (33, 34). The increased BRB dysfunction is associated with increased formation of CE and increased neuroinflammation. The role of C1q in BRB function is likely relevant in the context of FEVR and Norrie disease, related inherited diseases caused by impaired norrin/frizzled4 signaling. The genes mutated in these diseases include *NDP* (norrin) in Norrie disease and *FZD4*, *LRP5*, and *TSPAN12* in FEVR (41). Our data indicate that C1q maintains a degree of beta-catenin signaling in a FEVR mouse model, and that loss of C1q exacerbates BRB defects and neuroinflammation in this model. A role for C1q as a factor in BRB or BBB maintenance may extend to other retinal and neurological diseases beyond FEVR, including diabetic retinopathy and stroke.

Inherited deficiency of C1QA is a significant risk factor for SLE development (42), which may lead to lupus retinopathy (40). C1QA deficiency is thought to promote autoimmunity by impairing the clearance of apoptotic cells and nuclear antigens, which may cause the generation of autoantibodies. *C1qa* KO mice develop autoantibodies on specific genetic backgrounds, causing reduced long-term survival (35, 43). We performed our aging study on a C57BL/6J background that promotes survival but limits the development of autoimmunity. Potentially, the strong immunoglobulin extravasation in *Tspan12* KO^DBM^; *C1qa* KO compound mutant mice would promote autoimmune responses and lupus retinopathy with cotton-wool spots, and hemorrhages. The finding that these phenotypes were not present in *Tspan12* KO^DBM^; *C1qa* KO mice suggests that autoimmunity does not develop. Reasons include the genetic background, which may affect immune responses, or that cellular debris is still sufficiently cleared even in the absence of C1QA. *C1qa* deficient microglia may use alternative pathways to clear cellular debris. Potential alternative find-me signals, eat-me signals or opsonins include IgG, complement components C3b and C4b, pentraxins, or ApoE. Microglia may use Fc gamma receptors, complement receptors C3AR1 and C5AR1 or other receptors to detect these signals in a C1q-independent manner (44). Indeed, Fc gamma receptors, complement receptors, and complement components including C4b are strongly upregulated in *Tspan12 ECKO* with BRB dysfunction (7). Furthermore, the expansion of the microglia/monocyte-derived macrophage population may compensate for any reduction in the cellular rate of phagocytosis in *Tspan12* KO^DBM^; *C1qa* KO microglia/macrophages. Next to immune cells, astrocytes may phagocytose targets that are opsonized by C4b (39).

*Tspan12* KO^DBM^ and *Tspan12* KO^DBM^; *C1qa* KO compound mutant mice display a strong reduction of the ERG b-wave. The reduction of the ERG b-wave in treatment naïve conventional *Tspan12* KO mice, which are exposed to hypoxia in the inner nuclear layer due to angiogenesis defects (18), is only moderately more severe than the reduction of the ERG b-wave in *Tspan12* KO^DBM^ mice with normal angiogenesis and normoxia that we used in this study. While hypoxia likely contributes to the reduction of the ERG b-wave in conventional *Tspan12* KO mice, our findings highlight that BRB dysfunction also disturbs retinal homeostasis in a hypoxia-independent manner, for example by altering K^+^ ion homeostasis (45).

A second major conclusion from this study is that F4L5.13 agonist achieves complete resolution of CE in treatment naïve *Tspan12* ECKO mice. FZD4/LRP5 agonists also reduce neovascular tufts in mice with oxygen-induced retinopathy (16, 46); therefore, this class of agonists appears to target multiple processes relevant to diabetic retinopathy, including promoting BRB function and reducing pathological neovascularization. Our observation that F4L5.13 resolves CE further validates the norrin/frizzled4 signaling pathway as a compelling target for drug development in ME, for example in diabetic retinopathy, age-related macular degeneration, or retinal occlusive disease.

## Methods

*Sex as a biological variable.* Animals of both sexes were used for all studies. The study was not designed or powered to detect sex differences.

### Animals

*Tspan12* floxed (*Tspan12*^tm1.1Hjug^) and null (*Tspan12*^tm1.2Hjug^) alleles were reported previously (7). Tg(Cdh5-cre/ERT2)1Rha (47) was used as a EC-specific Cre driver and was provided by Dr. Ralf Adams under MTA with CANCERTOOLS.ORG. *C1qa* KO mice were obtained from Dr. Laura Nagy with permission from the owner of the material, Dr. Marina Botto/Imperial College. To ensure genetic consistency and survival of *C1qa* KO mice to the T_0_ + 12 months endpoint, all strains were backcrossed at least 7 generations with C57BL/6J mice. All mice were housed in a specific pathogen-free animal facility.

### F4L5.13 antibody administration

F4L5.13 (16) or vehicle (10 mM Histidine, 0.9% sucrose, 150 mM NaCl, pH 6.0) were administered intraperitoneally. *Tspan12* KO^DBM^ pups received 10 mg/kg at P6, P9, P12 and 4 mg/kg at P15, P18, P21, P24, P27. Adult mice received multiple doses of 10 mg/kg, i.p., as indicated in the respective figures.

### Fluorescein Angiography (FA)

Mice were anesthetized using an isoflurane gas delivery system. Immediately after anesthesia, pupils were dilated with a 1:1 mix of 1% tropicamide and 2.5% phenylephrine, administered as eyedrop. Systane Gel was applied to the cornea. 10 μl/g bodyweight of 0.25% fluorescein (diluted in sterile saline 0.9% from 10% Fluorescein-sodium, Akorn) was administered subcutaneously. Fluorescent fundus images were acquired 5 minutes after the fluorescein administration. A Micron III small animal fundus imaging system was used to acquire images. When FA was performed in the context of OCT, a Micron IV image guided OCT system was used (Phoenix Research Laboratories). Mice were placed on a heating pad during recovery from anesthesia.

### Image-guided Optical Coherence Tomography (OCT)

Mice were prepared for the experiment as described under FA. 10 OCT line scans were acquired, typically in the superior temporal quadrant as indicated by the red lines in FA images obtained during image-guided OCT. Line scans were acquired with a scan length of 1.45 mm and a spacing of 15 pixels. Images were captured using RevealOCT 2.1.6 software. Image processing (sharpening and contrast enhancement) was performed in Image J. CE was scored against the grading scale described in Supplemental Fig. S2 in a blinded fashion.

### Electroretinography

Electroretinograms were acquired under red light after overnight dark adaption. Mice were anesthetized using an isoflurane gas delivery system, pupils were dilated with a 1:1 mix of 1% tropicamide and 2.5% phenylephrine, administered as eyedrop, and hypromellose lubricant eye gel (Systane) was applied to both corneas. ERG recordings were obtained on a heated platform of a Celeris Diagnosys ERG system. The impedance ranged from 5 to 15 KΩ. The eyes received stimulation at 1 cd s/m2.

### Sulfo-NHS-LC-Biotin labeling

250 µl Sulfo-NHS-LC-Biotin (20 mg/ml) (Life Technologies, # 21335) was injected intraperitoneally. After 60 minutes, mice were anesthetized using an isoflurane drop jar, euthanized by cervical dislocation, and immediately transcardially perfused with 2 U/ml heparin in PBS.

### Wholemount retinal staining and quantification

Mouse eyes were dissected immediately after transcardial perfusion and mildly fixed in 4% PFA at RT for 15 minutes. After three washes with PBS, the retinas were dissected and blocked for 1 hour at RT with blocking buffer (5% goat serum, 0.5% Triton X-100 in PBS). Retinas were stained with streptavidin-Alexa 488 (Invitrogen, # S11223) or IBA-1 (Cell Signaling, # 17198) in blocking buffer overnight at 4°C. Retinas were washed 6 times (30 minutes/wash) at RT in PBS with 0.5% Triton X-100. IBA-1 staining was completed with secondary goat anti-rabbit-Alexa 488 (Life Technologies, # A11008,) in blocking buffer overnight at 4°C. Images were acquired using Keyence BZX800, and ImageJ was used for the quantification. For streptavidin quantification, 4x images were stitched. A threshold value was defined for the streptavidin-positive blood vessels of WT retinas, then this threshold value was used for all other groups to obtain the streptavidin-positive area above the threshold. For the quantification of IBA1-positive cells, image stacks were acquired at 20x magnification, and a maximum intensity projection was generated. IBA1-positive cells were manually counted from the projections.

### Quantitative PCR

RNA was extracted from mouse retinas using RNAzol-RT (ABP Bioscience, # FP314) according to the manufacturer’s instructions. Equal quantities of RNA were reverse-transcribed into cDNA using the Maxima First Strand cDNA Synthesis Kit (Thermo Fisher Scientific, # K-1642). Quantitative PCR (qPCR) was conducted with SYBR green detection and data analysis was performed using the ΔΔCt method. Primers for qPCR were designed to span exon-exon junctions. The following primers were used:

mGapdh:

Forward: 5’-GGGTGAGGCCGGTGCTGAGT-3’ Reverse: 5’-TCGGCAGAAGGGGCGGAGAT-3’

mTrem2:

Forward: 5’-CGAGGGTGCCCAAGTGGAAC-3’ Reverse: 5’-GGTGGTAGGCTAGAGGTGACCC-3’

mC4b:

Forward: 5’-GGCACACCTTGCCCGAAACA-3’ Reverse: 5’-AACCAAGCCCCAAAGGAGCC-3’

mAqp4:

Forward: 5’-GCTCGATCTTTTGGACCCGCA-3’

Reverse: 5’-GCACAGCGCCCATGATTGGT-3’

### Luciferase reporter assay and collection of serum

0.4 ml of 330 K/ml 293T cell suspension in high glucose DMEM (Corning, # 10-013-CV) with 5% mouse serum of the indicated genotype were seeded into each well of a 48-well plate. After 6 hours, cells were transfected with 160 ng DNA (4 ng FZD4, 8 ng LRP5, 8 ng GFP or TSPAN12, 140 ng reporter mix which was composed of TOPFlash plasmid, CMV-Renilla, and Lef1 as described (10)) using TransIT-LT1 (Mirus). 18 hours later, DualGlo luciferase assays were performed, and firefly and renilla luciferase signals were measured using a Synergy LX Multimode Reader (Agilent BioTek). The ratio of firefly/renilla luciferase signals was calculated, data were normalized to the reading obtained with WT mouse serum. To collect mouse serum, blood was withdrawn slowly via the left ventricle of deeply anesthetized mice using a 26-gauge needle. Mice were euthanized after blood collection. The blood samples sat undisturbed at room temperature for 30 minutes to allow clotting. Samples were spun at 2,000 x g for 10 minutes at RT. The clear supernatant (serum) was frozen.

### Statistics

A Shapiro-Wilk test for normality and Levene’s test for homogeneity of variance was performed. Parametric or non-parametric two-group and multi-group comparisons were performed as described in each figure legend. Parametric tests were homoscedastic or heteroscedastic T-tests or one-way ANOVA with Tukey post hoc, non-parametric tests were Mann-Whitney or Kruskal-Wallis tests. CE scores are on an ordinal scale, therefore these data were tested using a non-parametric test (48). Because cystoid edema and leakiness was markedly different between the two eyes of one mouse, we evaluated each retina separately without averaging. P<0.05 was considered significant.

### Study approval

All animal protocols were approved by the Animal Care and Use Committee of the University of Minnesota, Twin Cities.

### Data availability

The datasets analyzed for this study are available from the corresponding author upon reasonable request.

## Author contributions

LZ and HJJ wrote the manuscript. HJJ designed the study. LZ, JL, MA, HNJ, EO, KD, ET and HJJ conducted experiments. SM, HR, and ZC trained personnel or provided required instrumentation. SS and SA provided F4L5.13 antibody.

## Supporting information

Zhang et al. Supplememt

## Acknowledgements

This study was supported by grants from the NIH (R01EY024261 and R01EY033316 to HJJ and R21DA056728 to ZC), and from the Canadian Institute of Health Research (PJT-175160 to SA).

