## Supplementary material for "C1q limits cystoid edema by maintaining basal beta-catenin-dependent signaling and blood-retina barrier function": Zhang et al. Supplememt

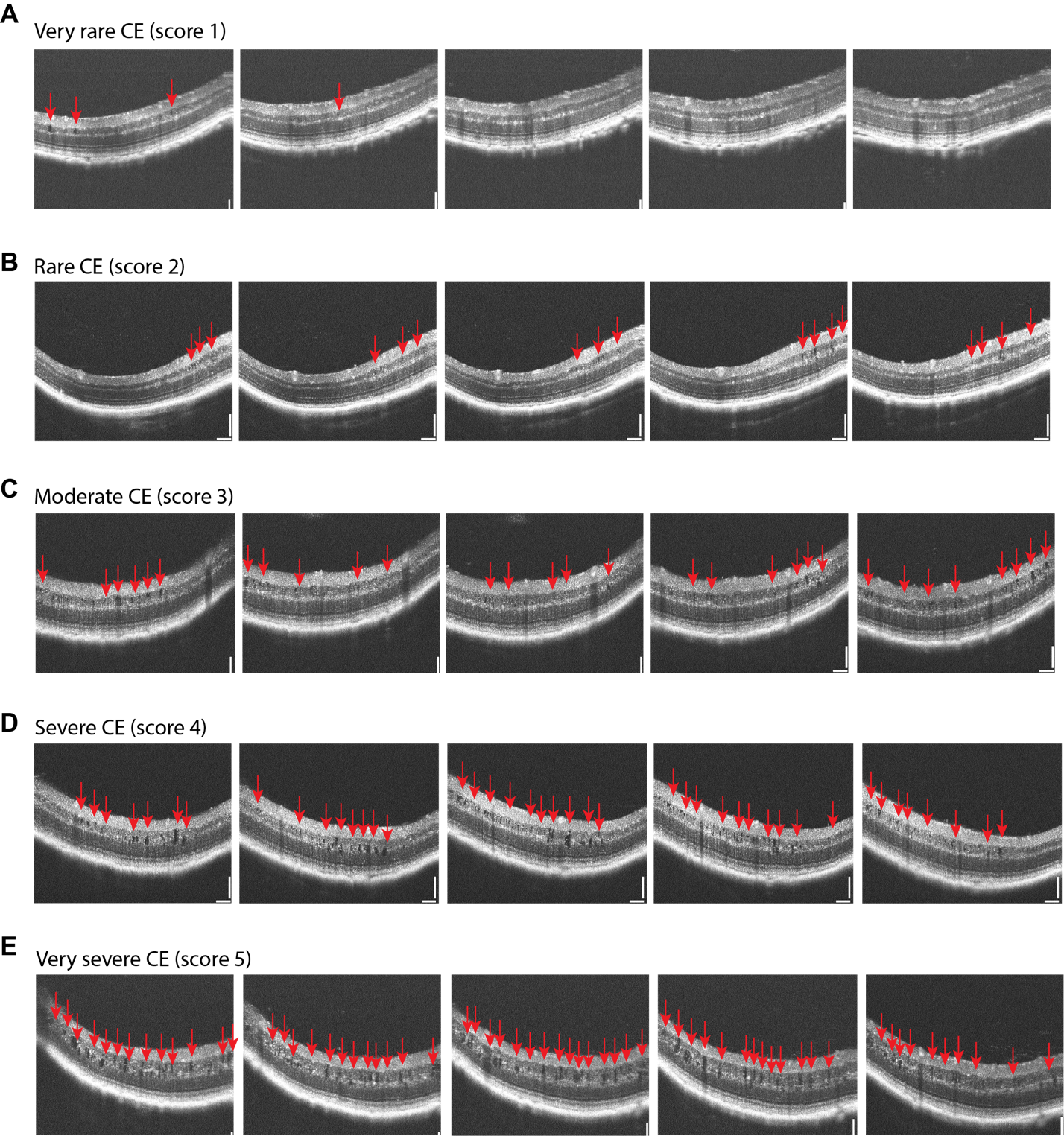

Supplemental Fig. S1. Representative OCT line scans representing CE of increasing severity, scored with 1 (very rare CE) to 5 (very severe CE). Five adjacent OCT line scans are shown per retina. Scale bars (100 µm) are not fully shown in all images due to cropping, the scale bars in panel B apply to all images.

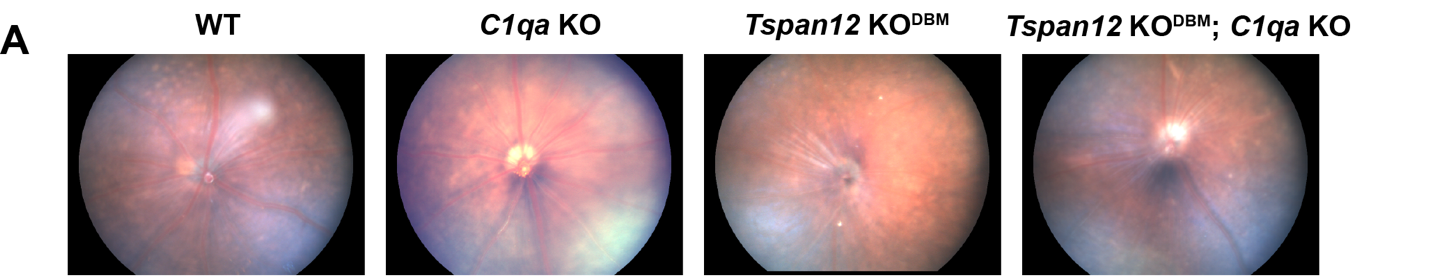

Supplemental Fig. S2. No hemorrhages or cottonwool spots in *Tspan12* KO^DBM^; *C1qa* KO mice. (A) Representative fundus images of N=3 mice per group. The images are focused onto the surface of the retina to allow detection of potential cotton-wool spots. SLE retinopathy was not detected in any of the four genotypes.
